# Uncovering perturbations in human hematopoiesis associated with healthy aging and myeloid malignancies at single cell resolution

**DOI:** 10.1101/2021.07.30.454542

**Authors:** Marina Ainciburu, Teresa Ezponda, Nerea Berastegui, Ana Alfonso-Pierola, Amaia Vilas-Zornoza, Patxi San Martin-Uriz, Diego Alignani, Jose Lamo de Espinosa, Mikel San Julian, Tamara Jimenez, Félix López, Sandra Muntion, Fermin Sanchez-Guijo, Antonieta Molero, Julia Montoro, Guillermo Serrano, Aintzane Diaz-Mazkiaran, Miren Lasaga, David Gomez-Cabrero, María Díez-Campelo, David Valcarcel, Mikel Hernaez, Juan P. Romero, Felipe Prosper

**Affiliations:** Area de Hemato-Oncología, Centro de Investigación Médica Aplicada (CIMA), Universidad de Navarra, IDISNA, Pamplona, Spain; Centro de Investigación Biomédica en Red de Cáncer, CIBERONC; Clínica Universidad de Navarra, Pamplona, Spain; Flow Cytometry Core, CIMA, Universidad de Navarra, Pamplona, Spain; Hospital Universitario de Salamanca-IBSAL, Salamanca, Spain; Department of Hematology, University Hospital Vall d’Hebron, Barcelona, Spain; Translational Bioinformatics Unit, NavarraBiomed, Pamplona, Spain; Computational Biology Program, CIMA, Pamplona, Spain; Red de Investigación Cooperativa en Terapia Celular TerCel, ISCIII

**Author notes:** contributed equally and should be considered joint first authors (Marina Ainciburu, Teresa Ezponda). contributed equally and should be considered joint senior authors (Juan P. Romero, Felipe Prosper). **Email addresses**, **Marina Ainciburu**, **Teresa Ezponda**, **Nerea Berastegui**, **Ana Alfonso-Pierola**, **Amaia Vilas-Zornoza**, **Patxi San Martin-Uriz**, **Diego Alignani** **Jose Lamo de Espinosa**, **Mikel San Julian**, **Tamara Jimenez**, **Félix López**, **Sandra Muntion**, **Fermín Sanchez-Guijo**, **Antonieta Molero**, **Julia Montoro**, **Guillermo Serrano**, **Aintzane Diaz-Mazkiaran**, **Miren Lasaga**, **David Gomez-Cabrero**, **María Díez-Campelo**, **David Valcarcel**, **Mikel Hernaez**, **Juan P. Romero**, **Felipe Prosper**.

## Abstract

Early hematopoiesis is a continuous process in which hematopoietic stem and progenitor cells (HSPCs) gradually differentiate toward specific lineages. Aging and myeloid malignant transformation are characterized by changes in the composition and regulation of HSPCs. In this study, we used single cell RNA sequencing (scRNAseq) to characterize an enriched population of human hematopoietic stem and progenitor cells (HSPC) obtained from young and elderly healthy individuals.

Based on their transcriptional profile, we identified changes in the proportions of progenitor compartments during aging, and differences in their functionality, as evidenced by gene set enrichment analysis. Trajectory inference revealed that altered gene expression dynamics accompanied cell differentiation, which could explain age-associated changes in hematopoiesis. Next, we focused on key regulators of transcription by constructing gene regulatory networks and detected regulons that were specifically active in elderly individuals. Using previous findings in healthy cells as a reference, we analyzed scRNA-seq data obtained from patients with myelodysplastic syndrome (MDS) and detected specific alterations of the expression dynamics of genes involved in erythroid differentiation in all patients with MDS such as TRIB2. In addition, the comparison between transcriptional programs and GRN regulating normal HSPCs and MDS HSPCs allowed identification of regulons that were specifically active in MDS cases such as SMAD1, HOXA6, POU2F2 and RUNX1 suggesting a role of these TF in the pathogenesis of the disease.

In summary, we demonstrate that the combination of single cell technologies with computational analysis tools enable the study of a variety of cellular mechanisms involved in early hematopoiesis and can be used to dissect perturbed differentiation trajectories associated with aging and malignant transformation. Furthermore, the identification of abnormal regulatory mechanisms associated with myeloid malignancies could be exploited for personalized therapeutic approaches in individual patients.

## INTRODUCTION

Mature blood and immune cells are generated by hematopoiesis, which is a well-characterized process that has been studied for more than a century ^(1)^. Classically, hematopoiesis has been modeled as a stepwise differentiation process, at the core of which reside hematopoietic stem cells (HSCs), which are common precursors with self-renewal capacity. A tree-like hierarchy arises from HSCs, in which lineage commitment occurs at binary branching points, giving rise to functionally and phenotypically homogeneous progenitor populations ^(2,3)^. However, recent single cell studies have questioned the validity of the classical model of hematopoiesis and have revealed heterogeneity within the HSC compartment and within progenitor populations that were previously considered homogeneous ^(4)^. Furthermore, it is difficult to establish boundaries between populations, which has led to the replacement of this concept by the idea of smooth transitions. Thus, early hematopoiesis is now viewed as a continuous landscape composed of undifferentiated hematopoietic stem and progenitor cells (HSPCs) with a variable degree of priming toward specific lineages (lymphoid, myeloid, or erythroid) ^(5–8)^.

During aging, the hematopoietic system undergoes various changes. There is evidence of an age-related increase in the relative number of HSCs, that presumably aims to compensate for a loss of repopulation ability ^(9,10)^. Despite this increase in the number of HSCs, the total bone marrow cellularity decreases ^(11)^. Additionally, a loss of lymphocyte production and a skewing toward myeloid differentiation have been demonstrated ^(12)^. Moreover, recent studies have revealed an increase in the number of platelet primed HSCs ^(13)^. A general loss of immune function that affects both innate and adaptive immunity has also been associated with aging ^(14)^. The molecular basis of these phenotypic changes includes an increased rate of random mutations in hematopoietic progenitors. Clonal hematopoiesis is a common trait of elderly individuals, in which blood cell subpopulations are clonally derived from a single HSC or progenitor with acquired mutations. Notably, some of these mutations are also associated with hematopoietic malignancies ^(15–18)^. Additionally, aging is also accompanied by transcriptional dysregulation. Gene expression analyses have revealed changes in cell cycle regulators ^(19)^, higher expression of myeloid signatures ^(13,19)^ and genes associated with leukemia ^(20)^, stronger response to inflammatory stimuli ^(21,22)^, and upregulation of pathways such as nuclear factor kappa beta (NF-κB) or tumor necrosis factor alpha (TNF-α) and downregulation of DNA repair ^(23)^. Lastly, epigenetic modifications that are in line with the transcriptional lesions detected have been observed in HSPCs obtained from elderly individuals ^(24)^.

Aging is associated with a higher risk of developing myeloid malignancies ^(25,26)^, suggesting a strong predisposition of hematopoietic cells from elderly individuals to lead to further alterations. Myelodysplastic syndromes (MDS) are among the main age-related hematological disorders. MDS are characterized by ineffective hematopoiesis and predisposition to transformation into acute myeloid leukemia ^(27)^, a highly aggressive neoplasm.

Collectively, previous evidence suggests the existence of a progressive decline in the hematopoietic system with age, which can result in disease if certain pathological events occur. HSC damage is the primary consequence of this process and its outcomes range from lineage bias to differentiation blockade and leukemic transformation ^(28)^. In this study, we examined these successive alterations in detail by considering hematopoiesis as a continuous process and analyzing the single cell transcriptional profile of early hematopoietic progenitors. We aimed to characterize key regulators of hematopoiesis in young and aged hematopoiesis and to apply this knowledge to identify specific perturbations occurring in patients with myelodysplastic syndromes. These regulators have the potential to predict, in a personalized manner, what abnormalities to expect and to identify potential therapeutic targets even at the early stages of transformation.

## RESULTS

### Transcriptional profiling of human young and elderly hematopoietic progenitor systems

To investigate the age-dependent changes in the hematopoietic system, we performed single-cell RNA sequencing (scRNA-seq) of bone marrow CD34^+^ cells obtained from five young (18-20-year-old) and three elderly (>65-year-old) healthy donors **(Figure 1a)**. Briefly, single-cell libraries were prepared using a Chromium controller instrument (10x Genomics) and sequenced up to an average depth of 30,000 reads per cell. A total of 34,590 and 40,641 cells were profiled from young and elderly donors, respectively. We first constructed an integrated dataset with cells from both young and elderly donors using an integration procedure implemented in the Seurat R package ^(29)^. Subsequently, we subjected the integrated dataset to quality control filtering and dimensional reduction to obtain a reference map for visualization (**Supplemental Table 1**). Secondary analyses including unsupervised clustering, identification of cluster markers, trajectory inference and gene regulatory network (GRN) reconstruction were performed independently for each group.

**Figure 1.**
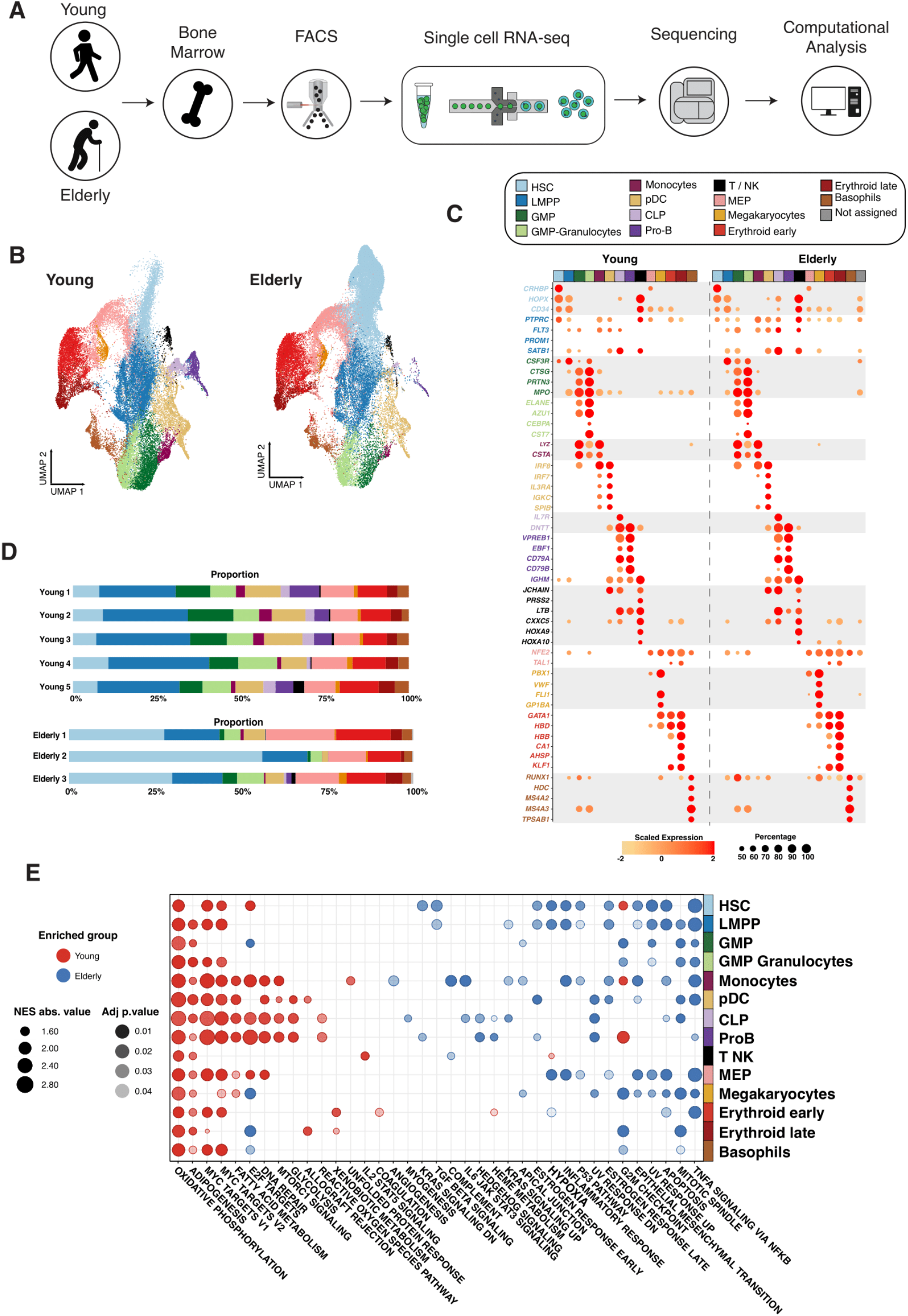
Transcriptional profiling of CD34^+^ cells from young and elderly healthy donors. a) CD34^+^ cells were obtained from bone marrow aspirates of young (n = 5) and elderly (n = 3) donors and subjected to single-cell RNA sequencing. b) UMAP plot with young cells colored according to unsupervised clustering results (left) and elderly cells labeled using an in-house cell classifier (right). c) Dot plot of cluster markers (adjusted *P*-value < 0.05) for the different cellular subpopulations identified. Dot size represents the percentage of cells that express each marker, and color represents scaled expression values. d) Bar plots showing the proportion of cells assigned to each cellular subpopulation for each donor independently. e) Dot plot of enriched terms after performing GSEA for each identified cluster. Dot color represents the enriched group, size indicates the NES absolute value, and transparency indicates the adjusted p value.

After integration, we extracted the cells obtained from young donors and subjected them to unsupervised clustering and differential expression analysis to identify cluster markers for manual cell type annotation and labeling (**Supplemental Table 2)**. This analysis identified 14 cellular subpopulations comprising the landscape of HSCs, early progenitors (MEP, LMPP, CLP, and GMP cells), and cells already committed to specific lineages (Pro-B cells, monocytic, erythroid, megakaryocytic, basophil and dendritic cell progenitors) (**Figure 1b).** Early progenitors were characterized by the expression of genes such as *CRHBP, HOPX* (initial lympho-myeloid differentiation), and *PBX1* (initial megakaryocyte and erythroid lineage). More mature progenitors presented myeloid (*MPO, CTSG, PRTN3, LYZ*, and *IRF8*), lymphoid (*DNTT, VPREB1*, and *EBF1*), erythroid (*HBD*, and *CA1*) and basophil (*HDC*, and *MS4A2*)-specific markers **(Figure 1c).** Once cellular subpopulations from the young donors were identified, we labeled cells of elderly individuals using a classification method based on a logistic regression with elastic-net regularization (see Materials and Methods). Regularized logistic regression has been previously applied to scRNA-seq data, due to its high interpretability and good performance with sparse input ^(30,31)^. We created binary classifiers that showed the probability that each cell belongs to a specific cluster **(Supplemental Figure 1a)** and assigned labels based on the highest probability. The performance of our classifier was determined by using a publicly available dataset containing 8,176 human CD34+ progenitors with known identities (Granja et al., 2019). This dataset was labeled using our method and Seurat **(Supplemental Figure 1b)** for benchmarking purposes. We obtained the proportion of cells classified within each cell type using both approaches **(Supplemental Figure 1c)**. Overall, both methods provided very similar results. Seurat displayed issues with early progenitors as 38% of HSCs were classified as MEPs, compared to 12% with GLMnet. This can also be observed within LMPPs and pDC cell types. The lymphoid lineage showed the biggest proportion changes however, this is related to the low number of B cells in the reference. As the GLMnet classifier performed better among progenitor cell types, we decided to use it in the rest of analyses.

When we applied the GLMnet classifier to the elderly individuals, we noted a significant increase in the proportion of HSCs and a reduction in the number of both committed lymphoid and myeloid lineages (Adj p.value < 0.05, **Supplemental Table 3**) as well as an increase in the proportion of erythroid-committed cells in the overall proportion, both in the integrated and individual datasets **(Figure 1d and Supplemental Figure 2).**

Next, we checked for functional differences between the young and elderly populations using gene set enrichment analysis (GSEA) after performing differential expression per cluster **(Supplemental Table 4)**. Cells from elderly donors exhibited a generalized enrichment of pathways activated in response to external molecules and insults, such as TNF-α, transforming growth factor beta (TGF-β), hypoxia or inflammation. Furthermore, the p53 pathway and apoptosis programs, which are activated by stressor stimuli, were also enriched in these cells **(Figure 1e)**. In particular, the TGF-β response and apoptosis genes, together with genes upregulated in response to ultraviolet radiation were specially enriched in age-associated HSCs. These results suggested the presence of an age-related response to the known more inflammatory microenvironment that is present in elderly individuals ^(32)^. Shifts in the abovementioned stress-related pathways were associated with increased expression in the elderly of potential drivers of such pathways, including *JUNB, FOSB, FOS* ID2 or *DDIT4* (**Supplemental Figure 3**). Hematopoietic progenitors from young donors were characterized by an increase in metabolism, including signatures related to glycolysis, fatty acid metabolism, and oxidative phosphorylation, in components of the respiratory chain (*UQCR, NDUF*, and *COX*), and the glycolytic enzymes *PGK1* and *TPI1*. These findings point towards an increased metabolic activity in progenitors from young donors. We also found that cell cycle progression (E2F targets) and proliferation (MYC targets) pathways were enriched in multiple cell populations from young donors **(Figure 1e)**, whereas *MYC* expression was generally downregulated in cells from elderly individuals, suggesting an age-mediated decreased proliferative activity. DNA repair was also found to be more active in cells isolated from young individuals. This goes in agreement with the known predisposition of age-associated progenitors to accumulate genetic lesions ^(33–35)^.

Overall, we found that the proportion of cell types changed with age, observing a decrease in the most mature lymphoid and myelomonocytic compartments. Furthermore, GSEA revealed age-related transcriptional alterations in all compartments, which suggests an altered biological behavior of HSPCs with age.

### Trajectory inference revealed age-specific drivers along lineages

To study the distinct differentiation trajectories observed during hematopoiesis and, specifically, alterations during aging, we subjected the single-cell expression datasets to trajectory inference using STREAM ^(36)^. We first inferred the differentiation trajectories of the young donors to obtain a reference that could serve as a basis for comparison. These trajectories revealed a common starting point that comprised early progenitors (HSCs, LMPPs, and MEPs). This initial root reached the branching point that divides cellular differentiation into the three main branches that resemble the main hematopoietic lineages: the myeloid, erythroid, and lymphoid lineages (**Figure 2a).** The identity of these branches was confirmed by plotting the expression profiles of previously described genes, such as *IRF8, GATA1*, and *EBF1* (**Figure 2b).** Then, in order to identify age-dependent alterations, the elderly dataset was projected onto the reference trajectories using the approach described in STREAM, (**Figure 2c**). We observed an increased proportion of HSCs in the root node, a reduced number of cells biased to both the myeloid and lymphoid lineages, and an increased number of progenitor cells committed to the erythroid compartment **(Figure 2d**).

**Figure 2.**
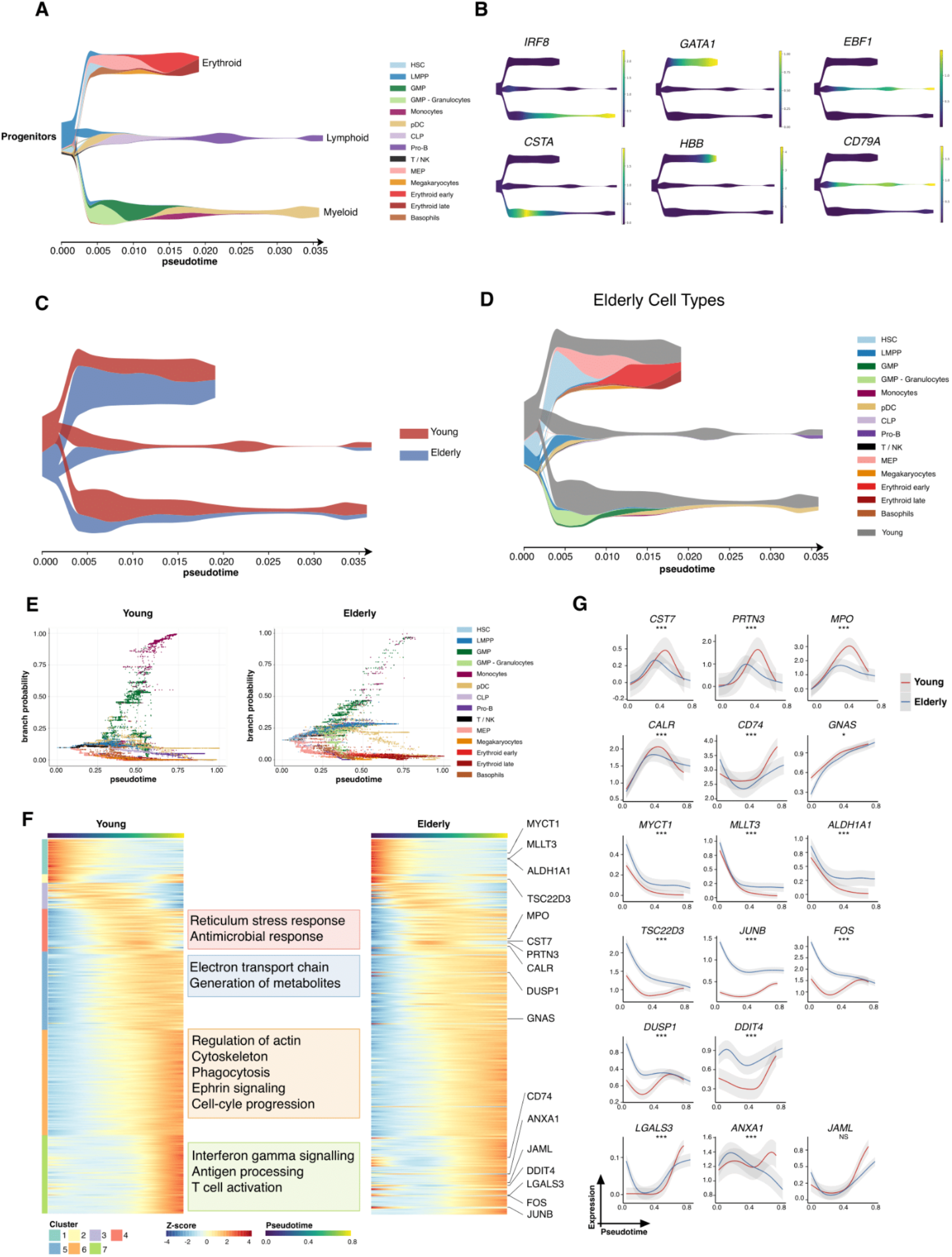
Trajectory inference of the hematopoietic lineages at single-cell resolution. a) STREAM plot obtained using cells from young healthy donors. Color denotes cellular subpopulations. The X-axis displays inferred pseudotime values. b) Expression of known cell-type markers for the different hematopoietic lineages projected in the STREAM plot. Color represents normalized expression values. c) STREAM plot of elderly differentiation trajectories projected in the young reference. Color indicates the proportion of cells belonging to each condition under study. d) STREAM plot of elderly differentiation trajectories projected in the young reference. Color represents the cell-type identity (gray color represents the proportion of young cells). e) Scatter plot of pseudotime vs. branch probabilities for the monocytic trajectory obtained using Palantir for young and elderly donors. Color represents the cellular subpopulation. f) Heatmap of gene expression trends for dynamic genes along the monocytic trajectory in young and elderly donors. The columns are arranged according to pseudotime values, and the rows are grouped according to gene clustering results. A summary of enriched terms for the gene clusters in young donors is shown. g) Expression trends in the comparison of young and elderly cells regarding the different genes involved in the monocytic trajectory. (NS = not significant, * adj p value < 0.05, ** adj p value < 0.01, *** adj p value < 0.001)

Next, we aimed to identify shifts in the gene expression dynamics along differentiation trajectories. We applied the Palantir algorithm ^(37)^, with which we recovered a general pseudotime (**Supplemental Figure 4a**) and differentiation potential (**Supplemental Figure 4b**) for each cell. We observed significant differences in the distributions of differentiation potential and pseudotime, with age-associated HSPCs accumulating greater differentiation potential and lower pseudotime (**Supplemental Figure 4d**). This suggested an immature cell state, which was consistent with our previous analysis. We could also define six trajectories corresponding to six committed compartments: i.e., erythroid, lymphoid, dendritic cells, monocytes, basophils, and megakaryocytes, and assigned each cell a probability of belonging to a specific branch **(Supplemental Figure 4c)**.

As an example of study of a particular trajectory, we focused on early monocytic differentiation, as it appeared to be impaired in the elderly donors. We plotted, in a per-cell basis, the associated pseudotime along the monocytic trajectory with the Palantir probability of reaching the final stage of the studied route (**Figure 2e**). This analysis revealed that HSCs and LMPPs from elderly donors had a higher probability of attaining the monocyte progenitor state, suggesting a stronger bias towards the monocytic compartment. However, GMPs displayed the opposite behavior which suggested that although age-associated progenitors appeared to have a stronger initial bias toward this lineage, a large number of more advanced progenitor cells lost the capacity for monocytic differentiation. This result is in line with the aged-associated decrease in monocytic progenitors described above.

To determine the transcriptional lesions that potentially alter monocytic differentiation during aging, we computed gene expression trends and clustered them based on their dynamics along the monocytic differentiation trajectory, following the Palantir pipeline. We characterized young gene clusters by overrepresentation analysis. Terms related to the immune system and monocytic function were enriched as the trajectory progressed, such as antimicrobial response or regulation of the actin cytoskeleton in intermediate stages, and antigen processing and presentation, or interferon gamma signaling, in gene clusters expressed near the terminal state. On computing trends for the cells of elderly donors, we observed that the overall behavior was similar. Nevertheless, genes with altered expression patterns in the elderly donors were visible (**Figue 2f**). Several myeloid differentiation markers (*CST7, PRTN3, MPO, CD74, CALR*, and *GNAS*) were expressed at lower levels across monocytic differentiation in the elderly donors, suggesting a less differentiated state. Accordingly, genes characteristic of stem or very early progenitor cells (*MYCT1*, *MLLT3*, and *ALDH1A1*) showed higher levels of expression in cells from the elderly donors across differentiation. Genes related to stress and inflammation response (*FOS, JUNB, TSC22D3, DUSP1*, and *DDIT4*), or monocyte chemotaxis and extravasation (*ANXA1, LGALS3*, and *JAML*), exhibited higher expression levels at the start of differentiation in the elderly, with several genes showing decreased expression at the terminal state (**Figure 2g**), further reinforcing the idea of a loss of capacity for monocytic differentiation in cells from elderly donors.

These analyses of the differentiation trajectory provided data regarding the genes that may be involved in the aberrant differentiation of age-associated HSCs. However, the regulation of these transcriptional programs cannot be inferred from this analysis.

### GRNs guiding young and elderly hematopoiesis

To elucidate the regulatory mechanisms underlying healthy hematopoietic differentiation, we constructed lineage-specific GRNs using SCENIC ^(38)^ for each of the datasets independently. Briefly, we obtained the set of activation regulons (transcription factor and their associated targets) for each of the cellular subpopulations and binarized its activity (on/off) in a per-cell basis using AUC values provided by SCENIC (through the AUCell algorithm). This approach enabled quantification of the proportion of cells that displayed an activated state for each regulon in each cluster. We then selected the top ranked regulons based on the regulon specificity score (RSS) **(Supplemental Figure 5a and 5b)** to identify specific regulatory mechanisms per cell type. For the young donors, we obtained a series of GRNs guided by transcription factors known to be relevant for specific cell populations, such as *HOXA9* in early progenitor cells, or *CEBPG, PAX5*, and *GATA1* in the myeloid, lymphoid and erythroid compartment, respectively (**Figure 3a)**. For some regulons AUC values were highly correlated with the corresponding gene expression profiles of the guiding TF, whereas others such as CEBPG or TCF3 displayed a more specific dynamism of GRN activity that is not observed with gene expression. For each of the hematopoietic lineages, we observed sets of regulons that regulated the transcriptional state of specific cellular differentiation programs and also the presence of common targets among different transcription factors (**Figure 3b**). This observation indicates that a single transcription factor might not only regulate cellular differentiation toward a specific route, but also contribute to other regulatory elements in other differentiation trajectories.

**Figure 3.**
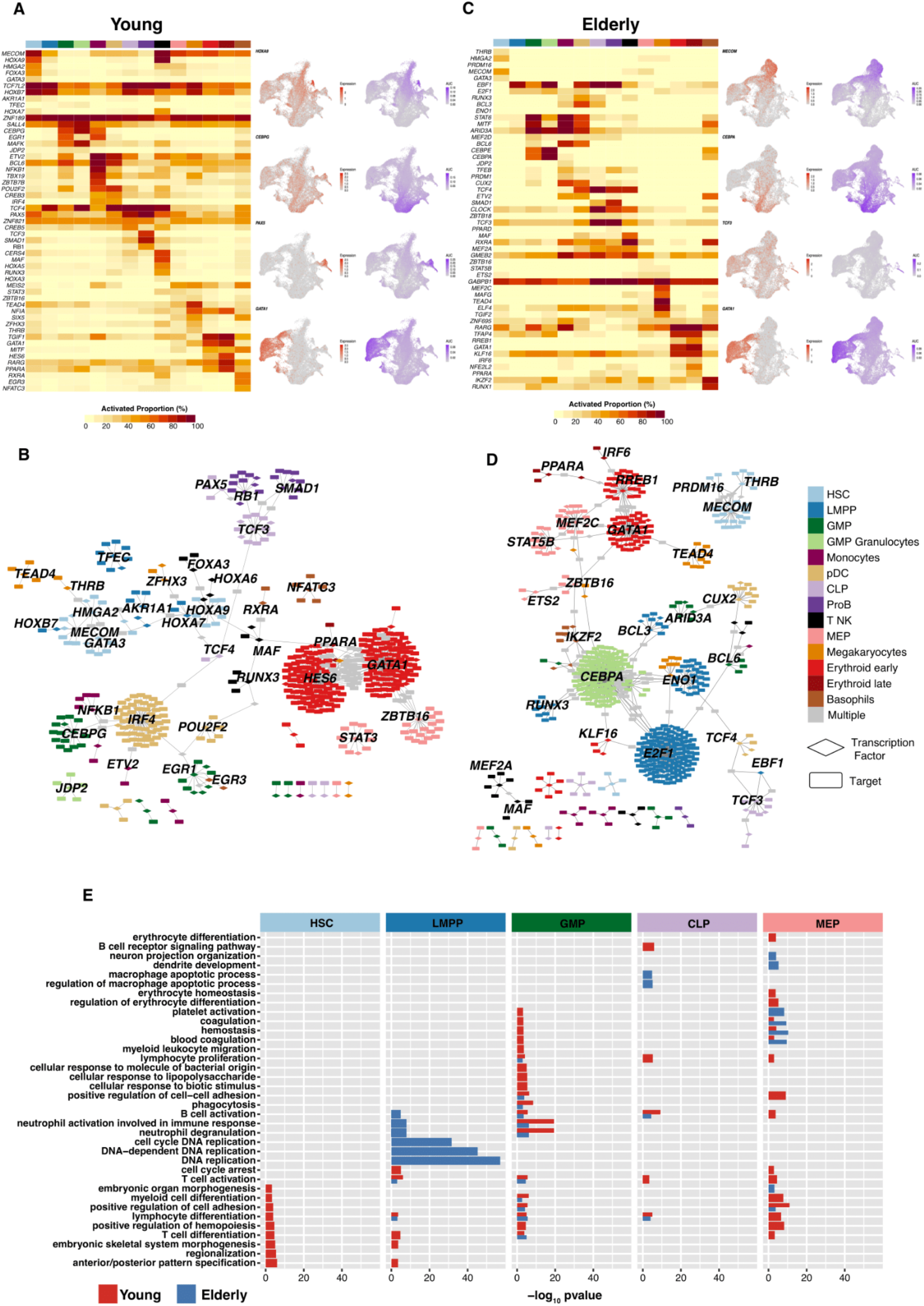
Gene regulatory network reconstruction of hematopoietic cellular populations. a) (Left) Heatmap showing the proportion of cells per cluster that have an activated state for different regulons in young cells. (Right) UMAP plots with normalized expression and AUC values for specific transcription factors. b) Gene regulatory network of the identified regulons for the hematopoietic system in young donors. Regulons were trimmed to include only targets with an importance score higher than the 3^rd^ quartile in each regulon. Node shape denotes gene-type identity, and color denotes cell population. Any target that can be assigned to multiple transcription factors is colored in gray. c) (Left) Heatmap showing the proportion of cells per cluster that have an activated state for different regulons in elderly cells. (Right) UMAP plots with normalized expression and AUC values for specific transcription factors. d) Gene regulatory network of the identified regulons for the hematopoietic system in elderly donors. Regulons were trimmed to include only the targets with an importance score higher than the 3^rd^ quantile in each regulon. Node shape denotes gene-type identity, and color denotes cell population. Any target that can be assigned to multiple transcription factors is colored in gray. e) Bar plot with enriched gene ontology categories after overrepresentation analysis. Categories are grouped per cell type, and color denotes the enriched group. Bar length represents statistical significance.

We performed the same analysis using the elderly dataset and observed that progenitor populations displayed a reduced proportion of cells with activated regulons. Specifically, HSCs showed a less active state of transcription factors, such as *MECOM* and *GATA3*, that were deemed specific to young donors (**Figure 3c)**. In already committed cellular subpopulations, we observed similar proportions of activated cells between young and elderly donors, with the erythroid lineage showing the most similar profiles. However, progenitors (GMP, MEP, and CLP cells) from elderly donors showed a lower number of cells with specific regulons activated. Overall, we observed reduced numbers of common targets **(Figure 3d)** between transcription factors, specifically for HSCs, which could result in the loss of co-regulatory mechanisms in the elderly donors.

We then performed gene ontology enrichment analyses using as input the genes composing the highest ranked regulons in early progenitor compartments (HSC, LMPP, GMP, CLP, and MEP cells). HSCs from young donors displayed enriched terms related to the differentiation of hematopoietic lineages, such as myeloid cell differentiation, lymphocyte cell differentiation and regulation of hematopoiesis, whereas HSCs from the elderly donors did not exhibit such enrichment, (**Figure 3e)**, suggesting a putative alteration of differentiation potential among aged HSCs. Although no increase in the percentage of LMPPs was observed in the elderly donors, this analysis indicated a clear enrichment in terms related to DNA replication in the elderly but not in the young donors, suggesting an aged-mediated alteration of the proliferative rate of this cell type. More mature progenitors also showed a youth-specific enrichment of differentiation processes such as lymphocyte differentiation, B-cell receptor signaling pathway, or T- and B-cell activation for CLPs, and erythrocyte differentiation and homeostasis for MEPs. This suggests that in age-associated progenitor cells, relevant regulators of hematopoiesis undergo functional alterations that result in the loss of expression of genes that are required for proper hematopoietic differentiation. Collectively, these results show aging-dependent alterations in the gene regulatory networks that guide hematopoiesis, which may be associated with the diminished differentiation capabilities that early progenitors present in the elderly.

### scRNA-seq analysis of MDS specimens reveals molecular lesions affecting normal hematopoietic differentiation

To further demonstrate the potential of the used computational methods in the prediction of the mechanisms underlying aberrant hematopoietic differentiation, we explored the transcriptional alterations that distinguish normal hematopoiesis during aging from abnormal hematopoiesis associated with the development of myelodysplastic syndromes. To this end, we performed scRNA-seq from bone marrow CD34^+^ cells obtained from four patients with MDS. In order to homogenize the group of study, and due to the great clinical and molecular heterogeneity of this disease, we focused our analyses in patients with a diagnosis of MDS with multilineage dysplasia.

We recovered 41,749 cells that passed the quality control filters (**Supplemental Table 5)**. The cell types were predicted using the GLMnet classification algorithm, as described previously (**Figure 4a**). We noted the absence of lymphoid compartments, considerable reduction in the number of HSCs, and increase in LMPPs and GMPs subpopulations compared to healthy elderly individuals. The total percentage of erythroid compartments varied among patients, although in general it was similar to that of normal counterparts (**Figure 4b**); nevertheless, the proportion of specific erythroid populations varied, with most MDS patients showing less MEPs and more late erythroid progenitors.

**Figure 4.**
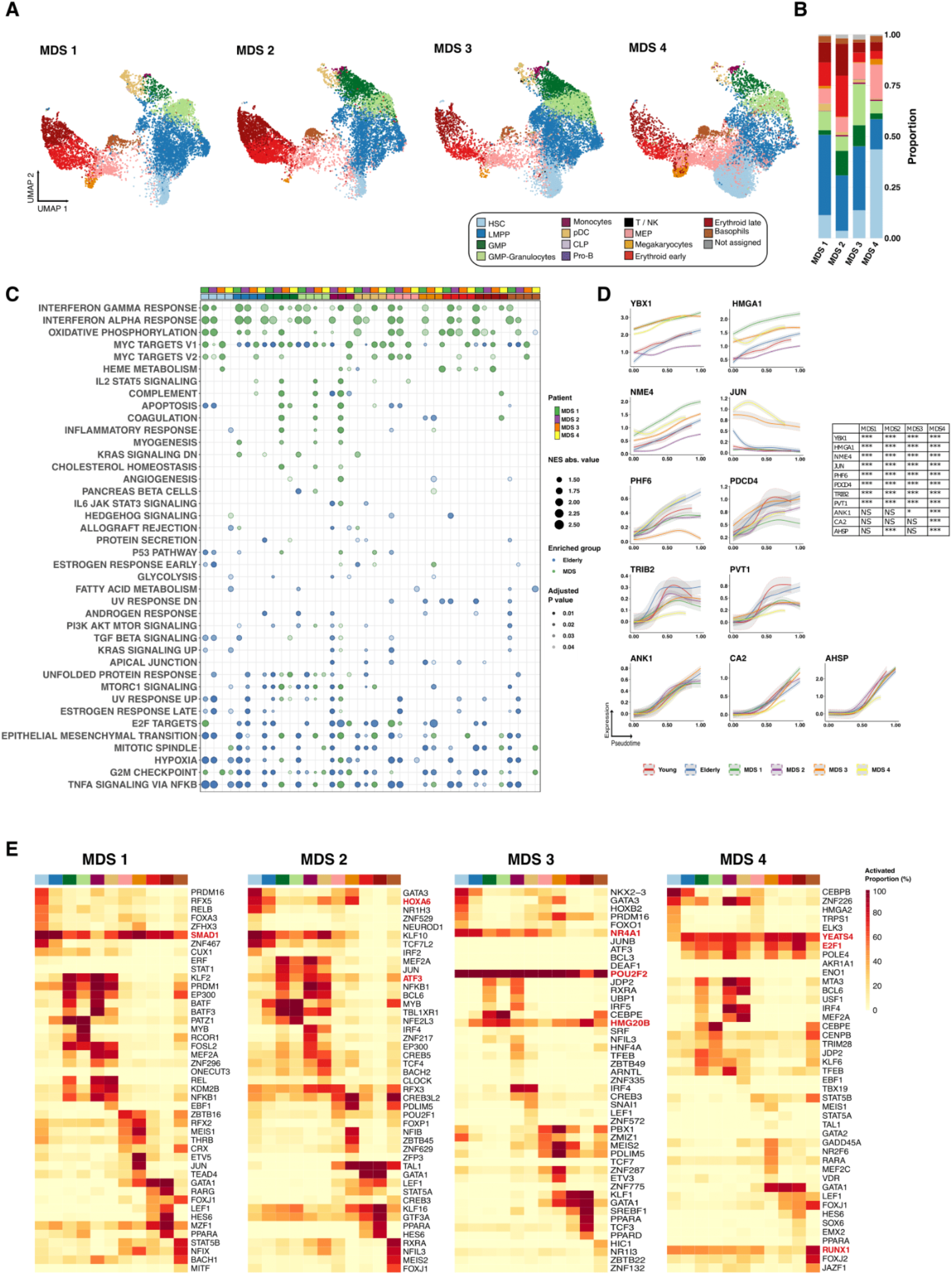
Computational analysis of pathological conditions, including myelodysplastic syndromess and acute myeloid leukemia. a) UMAP plot of CD34^+^ cells from MDS (n = 4). Cells are colored according to identity, as assessed using a previously described cell-type classification method. b) Bar plots showing the proportion of cells assigned to each cellular subpopulation for each donor independently. Color denotes the cellular subpopulation. c) GSEA results after performing differential expression between MDS and elderly donors. Dot color represents enrichment direction, transparency the statistical significance and size NES absolute value. d) Expression trends in the comparison of healthy and pathological cells regarding the different genes involved in the erythroid trajectory. (NS = not significant, * adj p value < 0.05, ** adj p value < 0.01, *** adj p value < 0.001) e) Heatmap showing the proportion of cells per cluster that had an activated state for different regulons in the 4 samples of patients with MDS among AML cells.

Next, we carried out GSEA of genes differentially expressed between healthy elderly donors and each of the patients for the detected subpopulations. Results showed potential aberrant functionality of MDS cells. The most evident alteration was the enrichment of MDS patients in genes related to interferon alpha and gamma response, which was more evident for cases MDS1 and MDS2. This observation is in accordance with previous reports demonstrating increased inflammatory signaling in the disease ^(39–41)^. Interestingly, genes associated with oxidative phosphorylation, a process that is very relevant for the metabolism of several types of cancer and that is considered as an emerging target ^(42)^, were more prominent in MDS. MDS patients also showed alterations of hallmarks related to cell proliferation, including E2F targets, mitotic spindle, or G2/M checkpoint, suggesting an aberrant proliferative activity of hematopoietic progenitors in these patients (**Figure 4c**). We also performed GSEA using the young donors as controls, and again found enrichment of interferon response in MDS patients. However, we observed variability in other potentially altered pathways, highlighting the importance of using age-matched controls for comparisons (**Supplemental Figure 6a**).

Most MDS patients are characterized by defects in erythropoiesis, showing dysplasia and/or citopenia of this lineage. Thus, as an example of the applicability of the trajectory inference analyses to identify MDS-related transcriptional alterations, we explored gene dynamics along the HSC-erythroid branch. We reconstructed the erythroid trajectory and computed gene trends along pseudotime using Palantir (**Supplemental Figure 6b**). Genes that were specifically expressed by any of the progenitors belonging to this branch (HSCs, MEPs, and erythroid progenitors) were selected and clustered according to their expression pattern (**Supplemental Figure 6c**). Although we did not detect broad changes in the transcriptional profiles among the conditions, we identified specific genes participating in erythroid differentiation that displayed different dynamics from both young and elderly healthy samples (**Figure 4d**). Examples of genes with altered dynamics included increased levels of genes with a negative role in erythropoiesis such as *JUN* and *YBX1* ^(43,44)^ and *NME4*, previously associated with poor prognosis in MDS ^(45)^. Moreover, we also observed decreased expression across the trajectory of factors that promote erythroid differentiation, such as *TRIB2, PHF6 and PDCD4* ^(46,47)^ and involved in erythroleukemia: PVT1 ^(48)^. Interestingly, whereas some of the genes showed aberrant dynamics in all the patients analyzed (i.e: *TRIB2*), most of them were altered in a subset or in individual patients, reinforcing the heterogeneity of MDS at the molecular level. Despite these changes, we observed that erythroid specific markers such as *ANK1, CA2* or *AHSP*, followed trends similar to those of healthy HSPCs, which is in agreement with the apparently normal progression of differentiation at the progenitor stages. These results indicate that, although at very early progenitor stages some drivers of erythroid differentiation exhibit normal trends of expression, others show clear abnormalities that may manifest in more mature stages as erythroid differentiation is altered and may be responsible for the anemia and/or erythroid dysplasia that characterizes MDS patients.

Using SCENIC we analyzed the transcriptional programs regulating HSPCs and reconstructed GRNs by extracting those regulons that were specifically active in each of the cellular compartments (**Supplemental table 6**). We identified active GRNs that were guided by TFs characteristic of the different populations (**Figure 4e** and **Supplemental Figure 6d**), and observed that a great number of the most prevalent regulons for each cell type were different from that of healthy donors. Interestingly, MDS cases showed regulons that were very active in most cell types, and that were not present in HSPCs from young or elderly samples, such as SMAD1 (MDS1); ATF3 and HOXA6 (MDS2); POU2F2, NR4A1, and HMG20B (MDS3); YEATS4, E2F1, and RUNX1 (MDS4). Interestingly, despite representing the same subtype, the individual MDS cases showed alterations of very specific regulons, demonstrating that the heterogeneity of the disease also takes place at the GRN level. Furthermore, some of the TFs guiding the regulons showing aberrant activity had previously been involved in the regulation of hematopoietic differentiation or the development of myeloid malignancies. For example, SMAD1 regulon was active in most cell types in patient MDS1, showing its highest activity in HSCs, LMPPs and erythroid precursors. SMAD1 knockdown has been shown to promote erythropoiesis, suggesting that high activity of this factor may negatively impact erythroid differentiation ^(49)^. Furthermore, SMAD1 pathway has been shown to be active in a model of persisting LSCs, suggesting that this factor may be relevant in the development of myeloid malignancies ^(50)^. Patient MDS2 demonstrated aberrant high activity of the regulon guided by ATF3, which has been shown to drive cell cycle progression in AML, and of that of HOXA6, a TF which potentiates hematopoietic cell differentiation and self-renewal. Patient MDS3 presented a ubiquitous activity of the regulon guided by POU2F2/OCT2, a TF overexpressed in AML, in all the subpopulations analyzed. This patient also showed high activity of NR4A1, a factor that has been shown to specify a distinct subpopulation of quiescent myeloid-biased HSCs ^(51)^, in HSCs and LMPPs. Moreover, we also detected that HMG20B, a known repressor of erythropoiesis ^(52)^, was prominently active in several cell types, including erythroid precursors. Finally, patient MDS4 demonstrated aberrant high activity of the regulons guided by YEATS4, which is amplified in different tumors, and E2F1, a TF whose increased activity has been previously described in MDS. These results suggested that aberrant activity of specific regulons in MDS patients could drive aberrant gene expression and ultimately promote a myelodysplastic phenotype.

Collectively, these data demonstrate the power of these analyses to identify novel transcriptional alterations in a personalized manner, and therefore to help uncover the molecular heterogeneity of the disease. Moreover, the approaches used do not only have the power to characterize genes with altered expression across MDS hematopoietic differentiation, but also to identify deregulated GRNs that could act as master regulators in the disease, which could be exploited therapeutically in individual patients.

## DISCUSSION

In this study we report the generation of high-resolution scRNA maps of human HSPCs and the changes associated with aging. Although previous studies focused on other species ^(22,53)^ and other layers of information such as mutations ^(33)^, proteomics ^(54)^, and proteo-genomics ^(55)^, our study represents one of the first analyses that describes early human hematopoiesis based on its dynamic gene expression and transcriptional regulation by identifying age-related changes that may be responsible for some of the phenotypic modifications observed during healthy aging. Applying this knowledge to HSPCs obtained from patients with MDS, we show how these analyses can help identify transcriptional alterations that play potential roles during disease development. This may ultimately lead to the identification of therapeutic targets.

Single-cell experiments are providing data at an unprecedented scale, which has allowed the identification of novel cell types and alterations that occur in biological systems. This increase in resolution has proceeded concurrently with the development of multiple computational tools that aspire to solve common problems arising with these technologies. An essential dilemma of this approach is how to establish reliable identities of the cells that will be used for subsequent analysis ^(56)^. In this study, we generated a reference system based on the transcriptional profile of HSPCs from healthy young donors and used it to assign labels to cells from elderly healthy and pathological donors. We generated an in-house cell classifier that allowed us to reduce the effect of technical artifacts that can contribute to erroneous prediction of cell identity. This classifier was applied to cells that underwent the same sorting, library preparation and sequencing procedures; thus, we expected to identify similar cell populations in each dataset. Additionally, when applied to external data, we found minor differences from the originally established labels, with the three lineages as well as the most immature states being precisely identified.

In accordance with previous studies, our analyses revealed an age-driven expansion of HSCs, which may represent a compensatory mechanism for less efficient hematopoiesis ^(9,10)^. We also noted a decrease in the proportion of lymphoid precursors, which is in agreement with the described aged-associated decrease in lymphoid differentiation capacity, as described previously ^(12)^. Using functional analyses, we also detected an enrichment of pathways related to apoptosis and inflammatory conditions in early progenitors among elderly donors, which can be explained by the inflammatory microenvironment known to be present in elderly individuals ^(32)^. Conversely, cells from young donors showed highly active differentiation and proliferation profiles, reinforcing the idea of the higher differentiation capacity among young individuals ^(28)^. In addition to these analyses, our study incorporated two other approaches: the characterization of transcriptional alterations across differentiation trajectories and the analysis of GRNs that guide different hematopoietic stages. Trajectory analyses pointed towards abnormal transcriptional dynamics across monocytic development during aging. We observed that although the most immature clusters (HSCs and LMPPs) showed an increased monocytic potential of differentiation in the elderly, more mature precursor cells presented reduced potential. These results would suggest that early immature cells are more primed toward the monocytic lineage during aging. However, alterations during later stages of differentiation result in a loss of monocytic differentiation capacity and, accordingly, a delay in the expression of genes that characterize the monocytic lineage. In agreement with our results, an age-associated skewing toward myeloid-biased HSCs and multipotent progenitor compartments has been described in mice ^(57)^. By focusing on GRNs, we observed that aged progenitor cell types showed a decrease in the activity of specific regulons, a lower degree of interaction between transcription factors and targets, and no enrichment of pathways involved in the cellular differentiation of specific lineages. Therefore, we were able to point toward regulatory factors guiding differentiation defects in the elderly.

We also shed light into the molecular pathogenesis of MDS. These syndromes are characterized by a significant phenotypic and genomic heterogeneity. In that context, our discoveries could have a direct clinical application by promoting the identification of patient-specific transcriptional lesions with a potential involvement in the differentiation defects. Although previous studies focused on the transcriptional lesions of MDS exist ^(58–64)^, they were performed using bulk populations of mononuclear cells and, thus, provide a limited perspective of the pathology in these patients. The detailed analysis of erythroid differentiation in our patients with MDS, led to the identification of genes with altered expression dynamics, which may play a key role in promoting dyserythropoiesis. Furthermore, GRN analysis revealed key regulons, whose activity could contribute to the phenotype of these cells. These results highlight the relevance of the methods used for understanding the mechanisms underlying differentiation defects in patients and provide new opportunities for the application of single-cell technologies in the clinic. This could help infer the transcriptional alterations involved in the aberrant differentiation of specific cell lineages in a patient-specific manner.

## MATERIALS AND METHODS

### Sample collection

The samples and data from the patients included in the study were provided by the Biobank of the University of Navarra and were processed according to standard operating procedures. Patients and healthy donors provided informed consent, and the study was approved by the Clinical Research Ethics Committee of the Clinica Universidad de Navarra. Bone marrow aspirates were obtained from healthy controls [young individuals (n= 5), median age, 20 years, range, 19-23 years] or patients undergoing orthopedic surgery [elderly donors (n=3), median age, 72 years, range, 61-84 years]. Samples from newly diagnosed patients with MDS were obtained from the Clinica Universidad de Navarra and collaborating hospitals. The patient’s clinical characteristics are shown in **Supplemental Table 5**.

### Fluorescence-activated cell sorting

For the CD34+ cells purification, bone marrow mononuclear cells (BMMCs) were isolated by Ficoll-Paque Plus (GE healthcare) density gradient centrifugation and stained using CD34 (clone 8G12; BD bioscience) CD64 (clone 10.1; Biolegend) CD19 (clone SJ25C1; Biolegend) CD10 (clone HI10A; Biolegend) CD3 (clone OKT3; BIolegend) CD36 (clone CLB-IVC7; Sanquin Plesmanlaan) CD61 (clone RUU-PL7F12; BD bioscience) for 15 min at RT. CD34+ CD64-CD19-CD10-CD3-CD36+CD61+ cells were then sorted in a BD FACSAria II (BD Biosciencs). Purified CD34+ cells were directly used for scRNA seq analysis.

### scRNA library preparation

The transcriptome of the bone marrow CD34^+^ cells were examined using NEXTGEM Single Cell 3′ Reagent Kits v3.1 (10× Genomics) according to the manufacturer’s instructions. Between 5,000 and 17,000 cells, depending on the donor, were loaded at a concentration of 700–1200 cells/μL onto a Chromium Controller instrument (10× Genomics) to generate single-cell gel bead-in-emulsions (GEMs). In this step, each cell was encapsulated with primers containing a fixed Illumina Read 1 sequence, a cell-identifying 16-bp 10× barcode, a 10-bp unique molecular identifier (UMI), and a poly-dT sequence. Upon cell lysis, reverse transcription yielded full-length, barcoded cDNA, which was then released from the GEMs, amplified using polymerase chain reaction, and purified using magnetic beads (SPRIselect, Beckman Coulter). Enzymatic fragmentation and size selection were used to optimize the cDNA size prior to library construction. Fragmented cDNA was then end-repaired, A-tailed, and ligated to Illumina adaptors. A final polymerase chain reaction amplification using barcoded primers was performed for sample indexing. Library quality control and quantification were performed using a Qubit 3.0 Fluorometer (Life Technologies) and an Agilent 4200 TapeStation System (Agilent), respectively. Sequencing was performed on a NextSeq500 instrument (Illumina) (Read1: 28 cycles; Read 55 cycles; i7 index: 8 cycles) at an average depth of 30,000 reads/cell.

### scRNA preprocessing

The demultiplexing of raw base call files (BCL) was performed using the 10× software cellranger mkfastq. The generated FASTQ files were aligned to the GRCh38 version of the human genome, and count matrices were constructed using cellranger count. The default barcode filtering was performed at this step. To remove doublets, we plotted the distribution of total genes and UMIs detected per cell and established a customized superior threshold for each sample (**Supplemental table 1**). Additionally, we filtered out cells with >5% or 10% of counts landing in mitochondrial genes, as this is an indicator of dying cells. In the elderly and pathological samples, we detected a small number of cells in which <1% of the mitochondrial genes clustered together; thus, we excluded them from the analysis.

### Data integration, clustering and visualization

Samples from the same condition (young, elderly or MDS) were integrated using the Seurat pipeline. Gene counts were divided by the total expression per cell, multiplied by a scaling factor of 10,000, and log transformed. Normalized counts were scaled across cells. The 2,000 genes with highest variance were selected using the variance stabilizing transformation method. Next, we performed integration as described previously (29) using 50 dimensions. After integration, we rescaled the data, regressed the cell cycle effect, and conducted principal component analysis (PCA). Based on the visual exploration of the scree plot, we selected the appropriate number of components to continue the analysis (**Supplemental table 1**).

Next, we performed unsupervised clustering on the young integrated data, using the algorithm implemented in Seurat. We constructed a shared nearest neighbor graph based on Euclidean distances in the PCA space with the chosen dimensionality and clustered the cells using the default Louvain algorithm. We tested several resolutions and assessed the results by calculating the average silhouette for each cluster. We determined the cluster markers using the Seurat function *FindAllMarkers* with the MAST method. Next, we annotated the clusters by manually inspecting the most specific markers and searching for curated markers in the literature.

To ease visualization, we embedded the cells from the young and elderly samples under the same coordinates. Thus, we integrated the young and elderly datasets using the pipeline described above. Using the first 30 principal components, we calculated the UMAP coordinates, as implemented in Seurat, which were then used in all the figures.

### Classification model

We constructed a classification model to predict the cell identity of elderly and MDS single cells using the previously annotated young dataset as a reference. To this end, we performed logistic regression with elastic-net regularization in the glmnet R package. This approach is appropriate for sparse input data, and the elastic-net penalty allows flexible selection of variables.

We constructed an individual binary prediction model for each of the 14 identities established. To train and validate each model, we selected all the cells from the appropriate cluster and an equal number of random cells from the remaining clusters. This set was then randomly divided in two: 75% was destined to training the model and 25% was dedicated to validation. We also performed an initial feature selection by selecting the subset of highly variable genes present in both the training–validation data (young) and the data to be classified (elderly, MDS, or AML). Initially, we trained models with different α values (1, 0.75, 0.5, 0.25, and 0.1). For each model, we used 10-fold cross validation, and the algorithm selected the optimal value for λ. Subsequently, we selected the most appropriate α value by evaluating the performance of the models on the validation dataset. If possible, we only retained the models that used between 20 and 150 variables. Among them, we selected the model with the maximum AUC and, in case of a tie, we successively searched for the minimum false-positive rate, minimum false-negative rate, and maximum number of variables. After the optimal model was selected, we used it to classify new data as positive or negative for a particular identity.

We repeated the steps from the creation of the training and validation sets to binary prediction, 10 times and stored the resulting predictions: both the class (positive or negative) and its associated probability of being positive. Finally, we averaged the 10 resulting probabilities for each binary model and assigned the identity that corresponded to the highest probability. In cases in which none of the identities were associated with a probability of >0.5, the cell was labeled as *not assigned.*

### GSEA

We performed differential expression analysis of cells belonging to the same identity to identify differences between conditions. We applied the Seurat *FindMarkers* function to the log-normalized counts using the MAST method and the options logfc.threshold = 0 and min.pct = 0. In order to control for batch effect, we set patients as latent variables. In addition, we tested for all genes expressed in every sample. Only cell types with at least 25 cells per condition were used. We ranked the genes according to their average log fold change. We performed GSEA using the fgsea R package and tested for hallmark gene sets.

### Trajectory analysis using STREAM

We extracted the differentiation branches in our data using STREAM. We then used the young batch-corrected matrix to establish the reference structure, onto which we then mapped the elderly cells. Using the STREAM pipeline, we computed the 2,000 most variable genes and 20 principal components. Then, we performed nonlinear dimensionality reduction using modified locally linear embedding. We projected cells onto a three-dimensional space and set the algorithm to use the 30 nearest neighbors. An elastic principal graph was constructed by computing 20 clusters and was refined using the following parameters: epg_alpha = 0.02, epg_mu = 0.05, and epg_lambda = 0.01. Subsequently, mapping was performed using the default STREAM parameters. For this step, we used the elderly batch-corrected matrix.

### Trajectory analysis using Palantir

Palantir was used to reconstruct hematopoietic lineages and recover gene expression dynamics along the lineages. This algorithm models differentiation and fate choice as a continuous probabilistic process. It orders cells along a global pseudotime, determines final states, and assigns each cell a probability to reach each terminal state.

We used the batch-corrected matrices to infer trajectories in young and elderly conditions. PCA was performed, and 30 diffusion components were calculated. For the young dataset, we selected 10 HSCs according to their position in the UMAP coordinates. We sequentially applied the Palantir algorithm using each of these cells as the initial state. Terminal states were stored for each run, and we retained those that appeared in more than five cases. If more than one terminal state corresponded to the same cell identity, we randomly selected one of them. We ran Palantir a final time adding the selected terminal states as prior information. For the elderly dataset, we selected the HSC with the highest probability, as determined by the GLMnet model, as the initial state. We then searched for cells with the highest similarity to the terminal states found in the young dataset. To this end, we calculated the nearest neighbor in the UMAP space. In the case of the MDS samples, we independently analyzed each patient. We used the normalized counts matrices, calculated 30 diffusion components, and set 10 eigen vectors to determine the multiscale space.

For each dataset, gene expression trends along pseudotime were calculated as described in Palantir using generalized additive models, with the addition of the branch probabilities as prior weights for the model.

### GRNs

Python implementation of SCENIC was used for GRN inference. All required inputs were downloaded from https://resources.aertslab.org/cistarget/ and https://pyscenic.readthedocs.io/en/latest/installation.html. Raw UMI matrices were provided as input for the pyscenic algorithm, as described in https://pyscenic.readthedocs.io/en/latest/tutorial.html. Regulon specificity scores were obtained using the *regulon_ specificity_ scores* command.

AUC matrices were imported into R and batch-corrected using the *removeBatchEffect* implemented in the limma R package. Batch-corrected AUC matrices were binarized using the BinarizeAUC function implemented in the AUCell R package. The percentage of active regulons per cell type was determined by counting the number of cells with a value of 1 in the binarized AUC matrix and dividing it by the number of cells per cell type. The top five regulons per cell type were selected according to the RSS scores.

Networks were drawn in Cytoscape, and only edges with an importance value larger than the 3^rd^ quartile (per regulon) were retained.

We performed over representation analysis for the different sets of genes that compose the top 5 specific regulons for HSC, LMPP, GMP, CLP and MEP, both in data from young and elderly donors. We tested for the gene ontology biological processes gene sets.

### Basic statistical analysis

We carried out pairwise comparison between pairs of proportions to test for significant differences in cell population proportions. We did so both among individuals from the same condition and between conditions. Two-samples wilcoxon test was performed to test for differences in Palantir pseudotime, differentiation potential and branch probabilities between young and elderly and per cell population. It was also used to find significant differences in gene expression trends across pseudotime between conditions. Multiple testing was addressed by adjusting p values using the Bonferroni-Holm correction method. Results were considered significant when adjusted p values < 0.05.

## Declarations

### Ethics approval and consent to participate

Samples and data from patients included in the study were provided by the Biobank of the University of Navarra and were processed following standard operating procedures approved by the Ethical and Scientific Committees.

### Consent for publication

Not applicable.

### Availability of data and material

All the single cell RNA sequencing data is available at Gene Expression Omnibus under accession number GSE180298

### Authors’ contributions

#### Conception and design

JPR, FP, MA, TE

#### Acquisition of data

NB, AAP, AVZ, PSMU, DA, JLE, MSJ, TJ, FL, SM, FSG, AM, JM

#### Computational analysis

MA, JPR, GS, ADM, ML, DGC, DV, MH

#### Writing of the manuscript

MA, TE, JPR,FP

#### Study supervision

JPR, FP

## Acknowledgements

We particularly acknowledge the patients for their participation and the Biobank of the University of Navarra for its collaboration.

## Additional Files

### Supplemental file 1

Supplemental figures S1 to S6

### Supplemental Table S1

Parameters used for scRNAseq analysis

### Supplemental Table S2

Cell type specific markers for each of the studied conditions (Adj p.value < 0.01 and logFC > 0.1)

### Supplemental Table S3

Cell type proportion test between young donors, elderly donors and conditions

### Supplemental Table S4

Differentially expressed genes between young and elderly per cell subpopulation (Adj p.value < 0.01 and |logFC| > 0.1)

### Supplemental Table S5

Clinical information from the donors and patients used in this study.

