## Supplemental Figure for "Uncovering perturbations in human hematopoiesis associated with healthy aging and myeloid malignancies at single cell resolution"

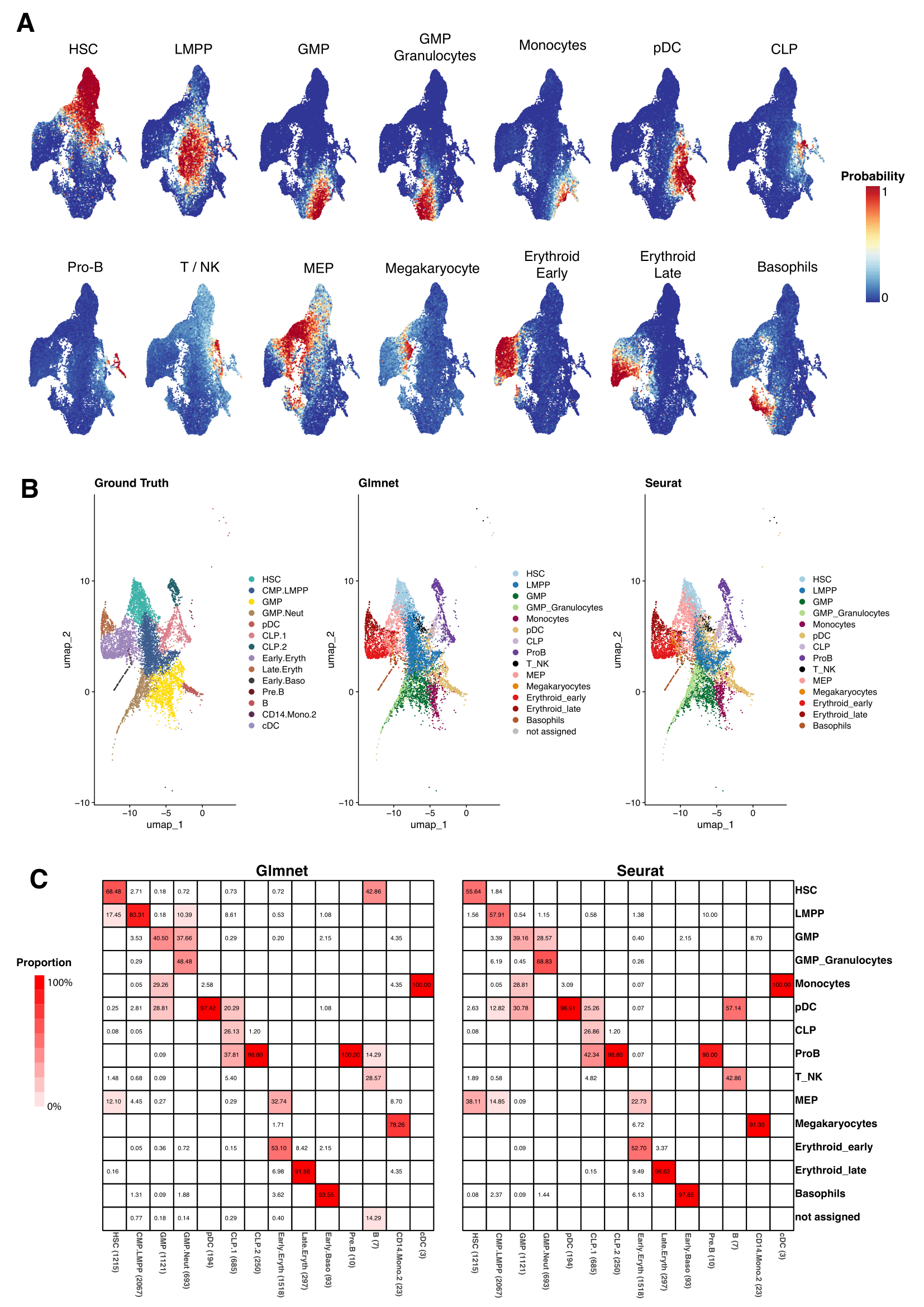

**Supplemental Figure 1.**

**Evaluation of GLMnet classification method A)** UMAP plots of the 3 elderly donors’ cells. They are colored by the probability of belonging to a specific identity, as computed on each of the binary classification models. **B)** CD34+ cells from Granja *et al* data. (left) cells are colored by the original classification. (middle) colored as the result of the GLMnet classification, using young donor data and identities as reference. (right) colored according to the predicted cellular identities using Seurat **C)** Heatmap showing the proportion of cells predicted within each of the ground truth groups. The sum per column equals to 100%.

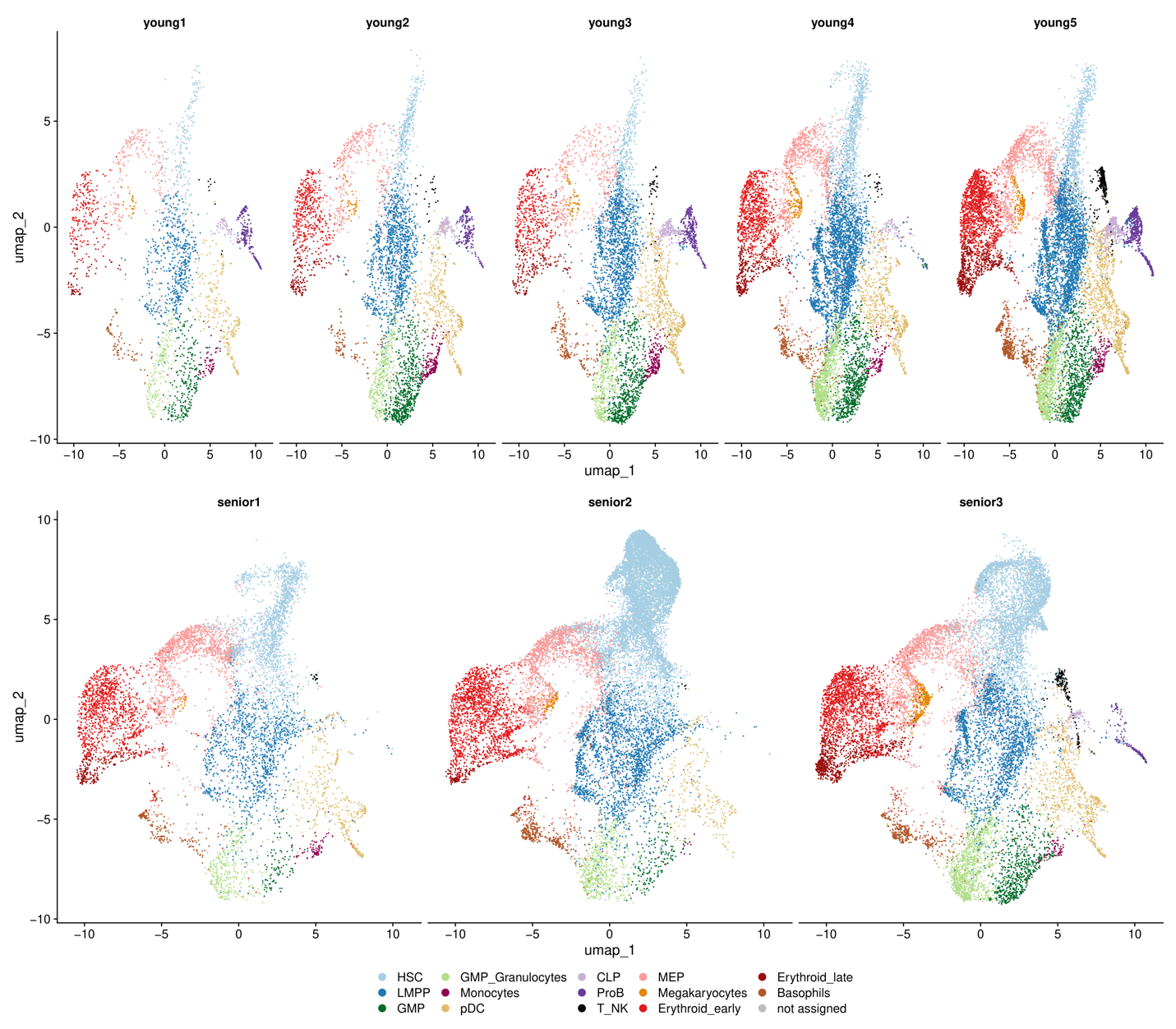
**Supplemental Figure 2.**

**Unsupervised clustering of CD34+ cells from young and elderly donors.** UMAP plot with cells colored by cellular subpopulation according to glmnet classifier. (top) young donors (bottom) elderly donors.

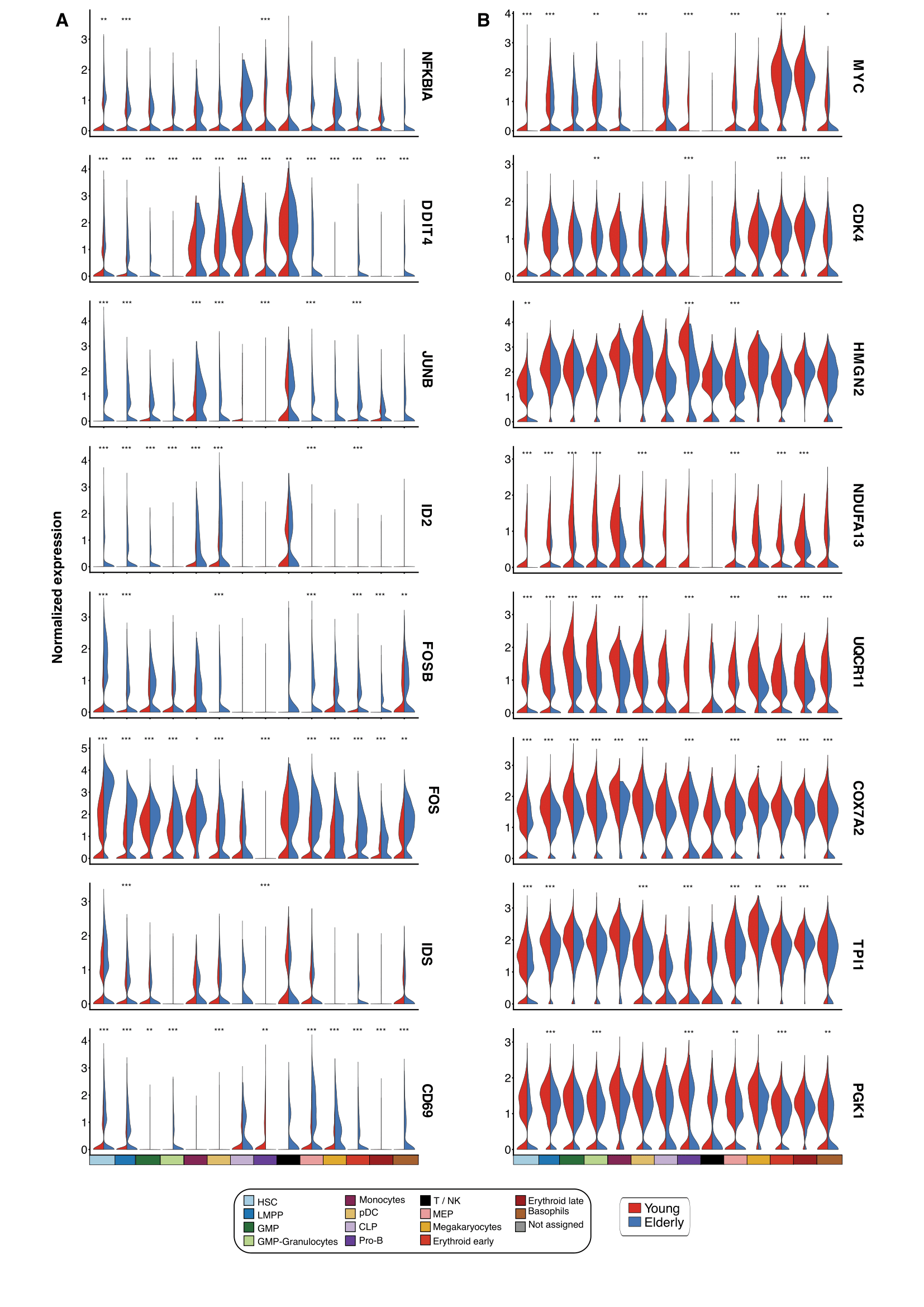

**Supplemental Figure 3.**

**Differentially expressed genes upon aging** Violin plots showing normalized expression of genes involved in differentially enriched pathways. Expression levels are divided by cell subpopulation and age (young cells colored in red and elderly cells in blue). **A)** Genes upregulated in elderly subpopulations **B)** Genes upregulated in young subpopulations. *adjusted p value < 0.05, ** adjusted p value < 0.01, ***adjusted p value < 0.001.

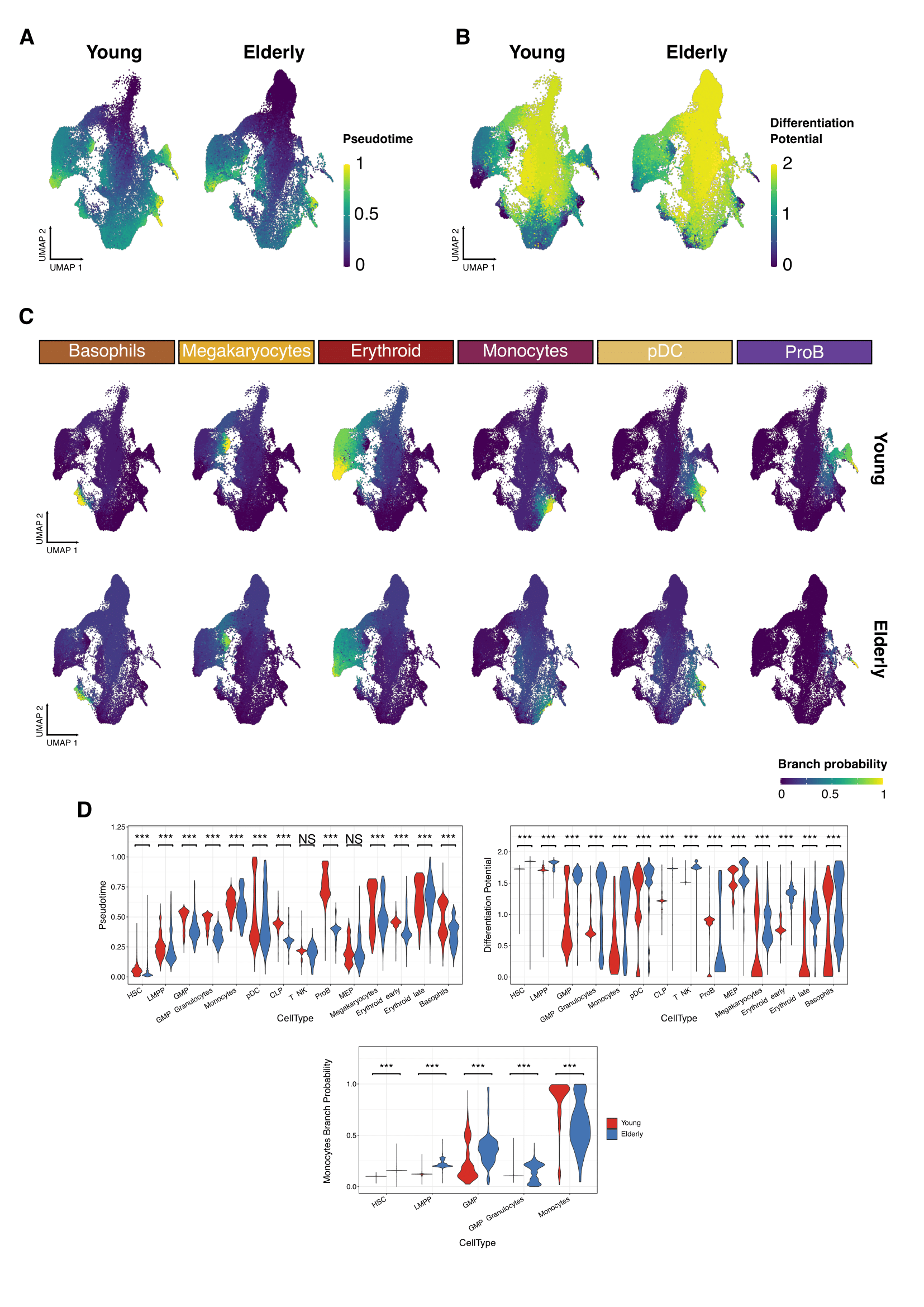

**Supplemental Figure 4.**

**Trajectory inference with Palantir reveals the main hematopoietic differentiation branches.** UMAP plots showing the results from applying Palantir algorithm to young and elderly cells. For both datasets, an HSC was established as initial state, based on UMAP coordinates. Final states were only indicated for the elderly dataset, as the UMAP nearest neighbors to the 6 young final points. Cells are colored by **A)** pseudotime **B)** differentiation potential **C)** Branch probabilities for each of the 6 differentiation paths retrieved. **D)** Violin plots colored by condition and representing the pseudotime and differentiation potential per cell type. (bottom) Branch probability for the differentiation route from HSCs to Monocytes. Wilcoxon two sample test *adjusted p value < 0.05, ** adjusted p value < 0.01, *** adjusted p value < 0.001, NS Non significant.

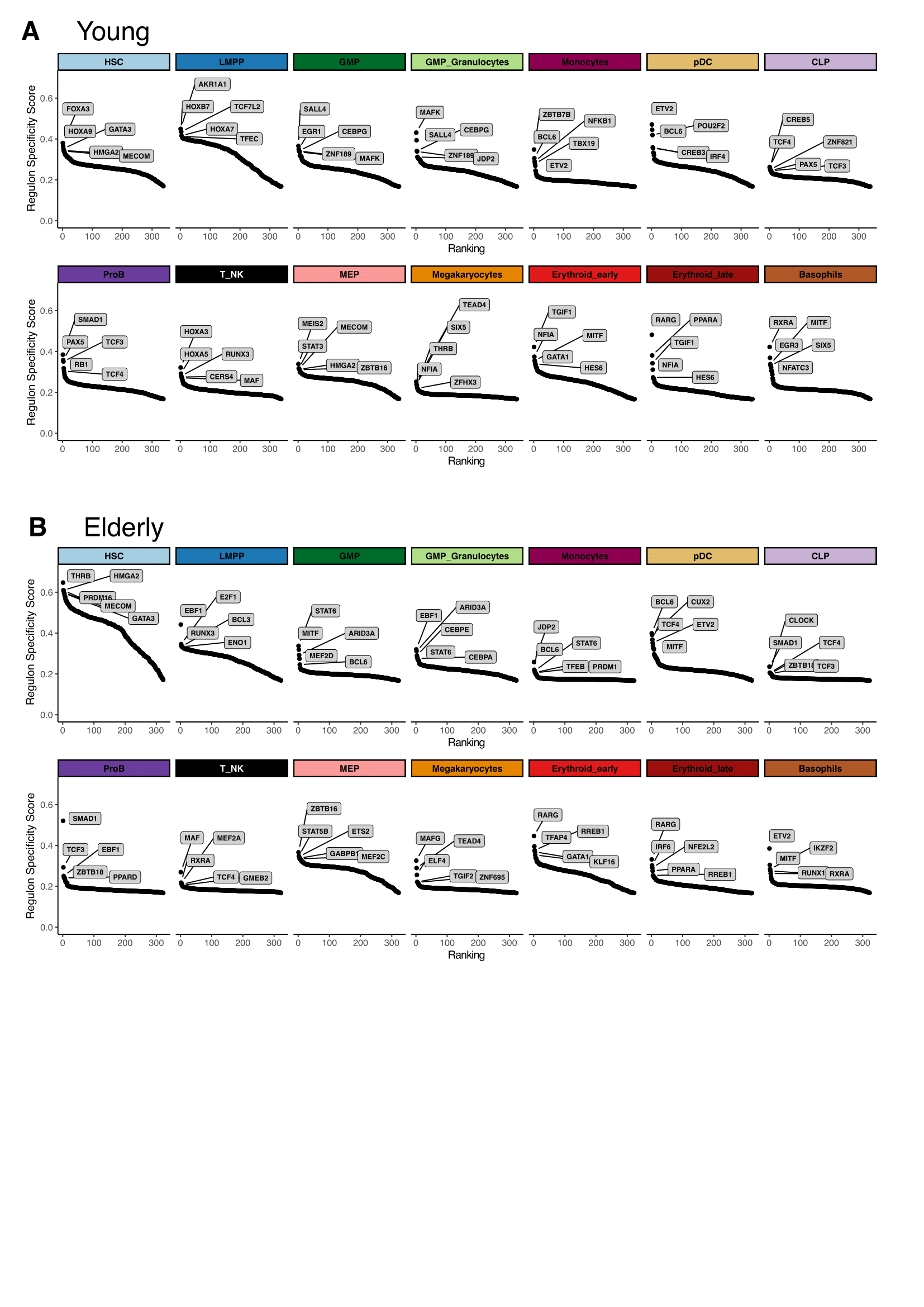
**Supplemental Figure 5.**

**Extraction of cell subpopulation specific regulons from Gene regulatory networks.** Regulons ranked by their specificity score (RSS), computed with pyscenic for each subpopulation. Names for the top 5 regulons with the most specific activity per subpopulation are shown. **A)** Young regulons **B)** Elderly regulons.

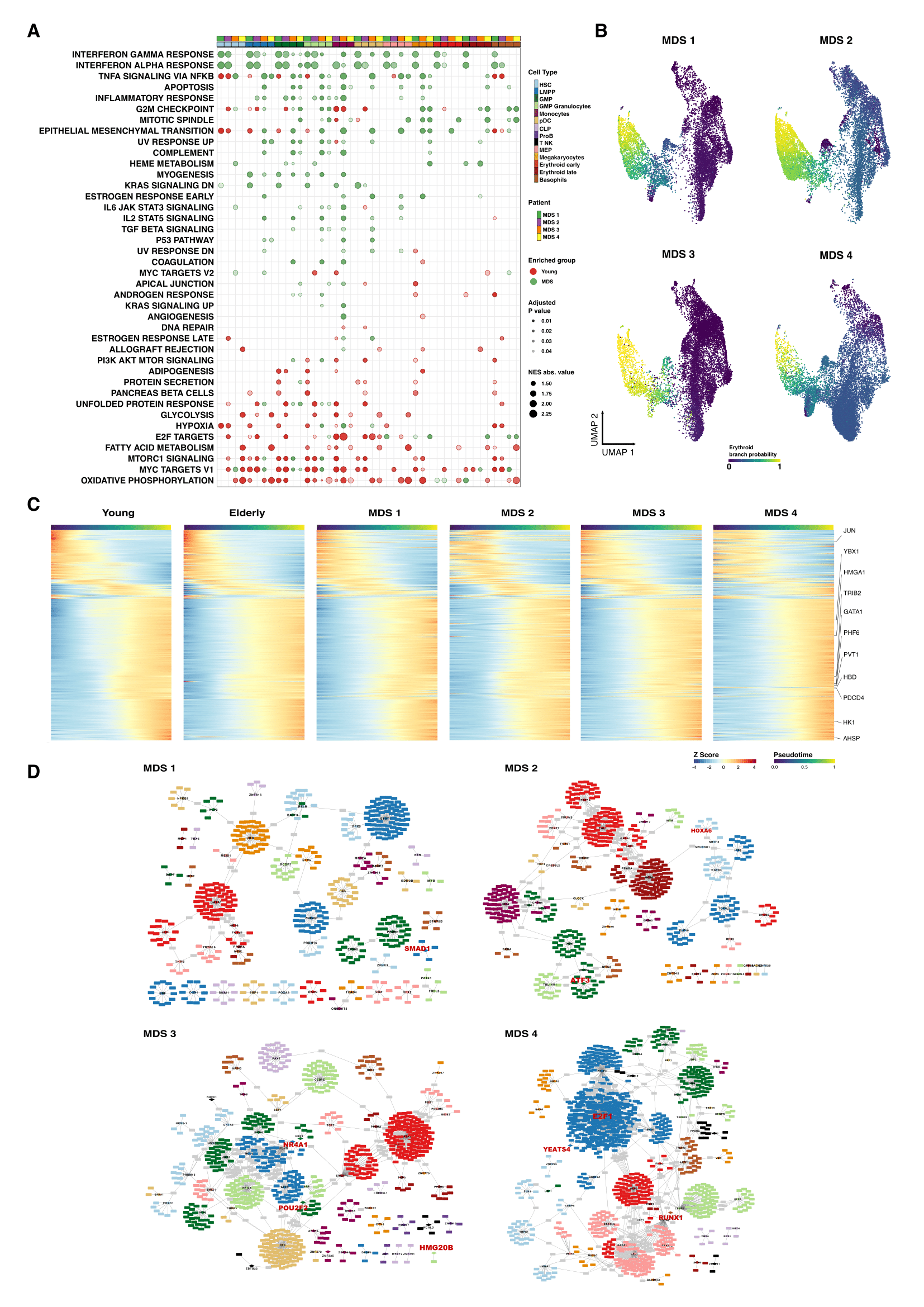
**Supplemental Figure 6.**

**Computational analysis of pathological samples. A)** GSEA results after performing differential expression between MDS and young donors. Dot color represents enrichment direction, transparency the statistical significance and size NES absolute value. **B)** UMAP with cells colored by Palantir probabilities for the erythroid trajectory. **C)** Heatmap of gene expression trends for dynamic genes along the erythroid trajectory in young, elderly and MDS donors **D**) Gene regulatory network of the identified regulons for MDS donors. Regulons were trimmed to include only the targets with an importance score higher than the 3rd quantile in each regulon. Node shape denotes gene-type identity, and color denotes cell population. Any target that can be assigned to multiple transcription factors is colored in gray. Important genes are labeled in red.
